# Structural Mechanism and Cellular Restriction of Tau Seeding from Endolysosomes

**DOI:** 10.64898/2026.06.08.731026

**Authors:** Eric Herrmann, Shuixia Tan, Kevin M. Rose, Richard M. Hooy, James H. Hurley

## Abstract

The prion-like spread of tau from cell to cell in the central nervous system involves escape from the endolysosomal network, which is counteracted by the lysosomal repair activity of the ESCRT system. Here, we investigate whether other components of the lysosomal damage sensing and repair system, namely the ESCRT-recruiting Ca^2+^ sensor ALG-2, conjugation of ATG8s to single membranes (CASM), the phosphoinositide-initiated tethering and lipid transport (PITT) pathway, and the Parkinson’s disease-related lipid transporter VPS13C are involved in tau spread. We found that the PITT pathway and VPS13C are strongly implicated in tau seeding by pre-formed fibrils (PFFs) in both neurons and astrocytes, CASM has a major role in astrocytes but not neurons, and ALG-2 has a lesser role in both. We then investigated the mechanism of damage and seeding by tau PFFs using cryo-electron tomography. Unlike the classical lysosome damage agent LLOMe, tau PFFs were not seen to directly interact with the lysosomal membrane, nor do they distort local membrane curvature. Lysosomes in PFF-treated cells were structurally intact. Extensive protein aggregates of similar character were seen in both the lysosomal lumen and in the cytosol proximal to lysosomes. The observations are consistent with the PFF-induced co-aggregation of tau with other cellular materials within lysosomes, with leakage to the cytosol attributed to reversible holes in the lysosome membrane.

## Introduction

The endolysosomal network (ELN) is the cell’s digestive tract. The ELN is responsible for processing both inbound endocytic materials and cellular contents taken up by autophagy. In the course of its function, the ELN is exposed to a vast range of potentially damaging materials, including infectious pathogens and amyloid aggregates^1,2^. The importance of ELN degradative function is highlighted by the ubiquity of mutations in ELN-related genes in neurodegenerative diseases, including Parkinson’s disease (PD), Alzheimer’s disease (AD), amyotrophic lateral sclerosis (ALS), and frontotemporal degeneration (FTD)^3,4^.

Many neurodegenerative conditions, most notably PD and AD, are driven by the abnormal misfolding and aggregation of endogenous proteins into cross-β amyloid structures^5^. A defining characteristic of AD and related tauopathies is the progressive cell-to-cell spread of tau, a microtubule-associated protein, throughout the brain^6^. This progressive spread is increasingly understood to operate via a prion-like mechanism^7^. Pathological tau seeds are released from donor cells and taken up by recipient cells via endocytosis^8^. Once internalized, these seeds physically interact with endogenous tau, nucleating the polymerization of cytosolic tau and driving disease progression.

For this pathogenic templating to occur, internalized tau seeds must escape the confinement of the endolysosomal system to reach the cytosol. The repair and degradative capacity of the ELN has a direct bearing on the prion-like spread of tau aggregates in AD^9^. In the current leading model for this process, tau spread relies on the ability of endocytosed tau seeds to escape vesicular confinement and seed the aggregation of cytosolic tau^10^. Cellular membrane repair responses exist that can in principle resist membrane permeation and so resist the escape of lumenal materials^11^. The ESCRT machinery, long known for its role in membrane budding and sealing, is a primary responder to nanoscale lysosomal permeabilization^12^. ESCRTs have been shown to counteract tau spread^13,14^. The ESCRTs cooperate with other lysosomal damage repair mechanisms. ESCRTs response to Ca^2+^ release as sensed by ALG-2 and likely other Ca^2+^ -binding proteins^11,12,15^. Conjugation of ATG8s to single membranes (CASM) signals damage^16^, while VPS13C^17^ and the phosphoinositide-initiated tethering and lipid transport (PITT) pathway^18^ source phospholipids from the ER to replace lipid molecules lost to damage. The roles in tau spread of ALG-2, CASM, VPS13C, and PITT, are, however, unknown. One goal of this study was to understand whether some or all of these interrelated repair systems counteract tau seeding in key cell types of the central nervous system (CNS), astrocytes and neurons.

At the molecular level, interrogating this defensive network requires targeting its specific components. The localized calcium efflux caused by permeabilization is sensed by the penta-EF-hand protein ALG-2, which recruits the ESCRT-III machinery for repair^11,13^. The CASM pathway can by triggered by ATG16L1 or TECPR1^19–21^ and their shared cofactor ATG5-ATG12^1,2^ Concurrently, structural membrane patching via the PITT pathway is initiated by the lipid kinase PI4K2A^18,22^ and executed by bulk lipid transfer proteins (BLTPs), including VPS13C^23^ and the autophagy and PITT factors ATG2A and ATG2B^18^. Because ATG2A and ATG2B are functionally redundant, a double knockdown (DKO) is needed to completely ablate ATG2 activity^24,25^.

We employed a GFP-K18 tau reporter system^8,13^ to monitor PFF-induced tau aggregation in human brain-derived astrocytes and iPSC derived iNeurons. By systematically knocking down key components of the membrane damage response pathways in primary astrocytes and iNeurons, we assessed their effects on tau seeding. We then used *in situ* cryo-electron tomography (cryo-ET) to directly visualize the compromised lysosomes and tau aggregation. By integrating these functional and structural data, we provide a refined molecular model detailing how cellular containment ultimately fails, allowing tau to seed aggregation, which spreads into the cytosol.

## Results

### CASM restricts tau spread in astrocytes but not neurons

To investigate the mechanisms restricting tau escape, we utilized a K18-GFP reporter system^8,13^ alongside the lysosomal marker TMEM192-RFP (Figure S1a). In human iPSC-derived iNeurons, incubation with tau PFFs induced the aggregation of K18-GFP into fluorescent puncta (Figure S1b-f), as demonstrated previously in HeLa cells and astrocytes^8,26^. To interrogate the specific pathways involved, we generated lentiCRISPR-mediated knockdowns of key components of the ESCRT recruitment system, CASM, and the PITT pathway (Figure S2).

The role of calcium leakage responses was determined by first knocked down the calcium sensor ALG-2 in astrocytes co-expressing K18-GFP and TMEM192-RFP (Figure S2a). Following tau PFF treatment, astrocytes depleted of ALG-2 (Figure 1b) exhibited a significant increase in the ratio of K18-GFP puncta-positive cells compared to controls. To investigate the role of CASM factors, K18-GFP/TMEM192-RFP-expressing astrocytes were infected with lentiCRISPR viruses against TECPR1 and ATG12. Following PFF treatment, the depletion of either TECPR1 (Figure 1c) or ATG12 (Figure 1d) increased the ratio of K18-GFP puncta-positive astrocytes.

**Figure 1:**
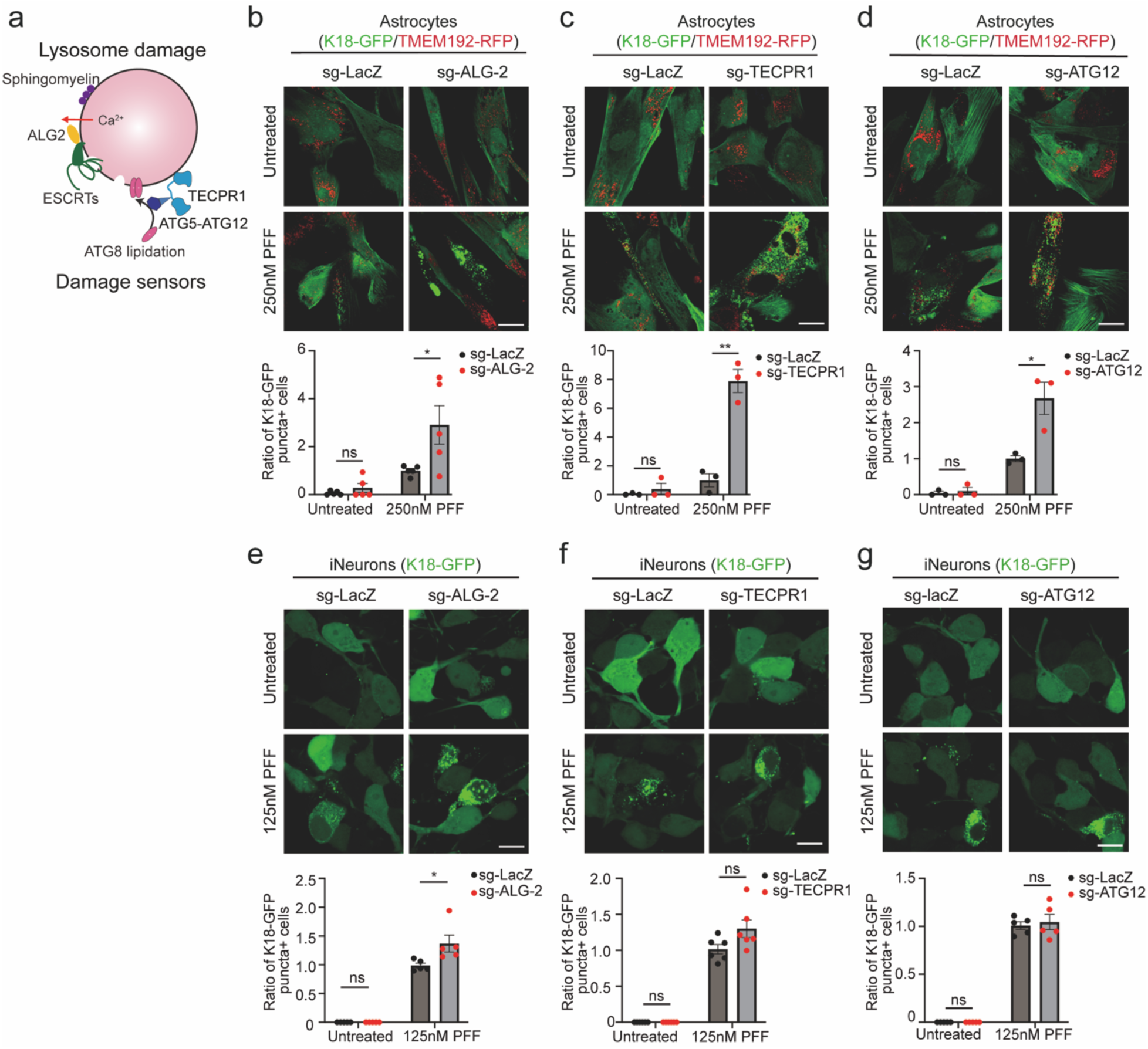
ALG-2 and CASM as lysosomal damage sensors prevent K18.Tau aggregation. (a). Schematic representation of ALG-2 and CASM sensing lysosomal damage and initiating membrane repair. (b). K18-GFP/TMEM192-RFP-expressing astrocytes were infected with sg-ALG-2 lentiCRISPR virus, followed by Tau PFF treatment. Then the ratio of K18-GFP puncta-positive cells was quantified (n = 5 independent experiments, 463-752 cells per group were quantified). (c). K18-GFP/TMEM192-RFP-expressing astrocytes were infected with sg-TECPR1 lentiCRISPR virus, followed by Tau PFF treatment. Then the ratio of K18-GFP puncta-positive cells was quantified (n = 3 independent experiments, 394-522 cells per group were quantified). (d). K18-GFP/TMEM192-RFP-expressing astrocytes were infected with sg-ATG12 lentiCRISPR virus, followed by Tau PFF treatment. Then the ratio of K18-GFP puncta-positive cells was quantified (n = 3 independent experiments, 302-403 cells per group were quantified). (e). iNeurons were infected with sg-ALG-2 lentiCRISPR virus to achieve ALG-2 knockdown, followed by K18-GFP lentivirus infection and tau PFF treatment. Then the relative ratio of K18-GFP puncta-positive cells was quantified (n = 5 independent experiments, 226-493 cells per group were quantified). (f). iNeurons were infected with sg-TECPR1 lentiCRISPR virus to achieve TECPR1 knockdown, followed by K18-GFP lentivirus infection and tau PFF treatment. Then the relative ratio of K18-GFP puncta-positive cells was quantified (n = 6 independent experiments, 509-715 cells per group were quantified). (g). iNeurons were infected with sg-ATG12 lentiCRISPR virus to achieve ATG12 knockdown, followed by K18-GFP lentivirus infection and tau PFF treatment. Then the relative ratio of K18-GFP puncta-positive cells was quantified (n = 5 independent experiments, 273-399 cells per group were quantified). The ratio of K18-GFP puncta+ cells shown in the histogram was normalized to that of the Tau PFF-treated sg-LacZ group. An unpaired two-tailed Student’s t-test was used for pairwise comparisons. Scale bars were 20μm, and 10μm in iNeurons.

To determine if the damage sensing responses mediated by ALG-2 and CASM are conserved in neurons, we next evaluated ALG-2, TECPR1 and ATG12 depletion in the iNeuron model. Consistent with the astrocyte data, knockdown of ALG-2 resulted in a significant increase in the relative ratio of K18-GFP puncta-positive cells upon PFF exposure (Figure 1e). However, the knockdowns of TECPR1 and ATG12 do not have a significant effect on the relative ratio of K18-GFP puncta (Figure 1f,g).

### PITT and BLTPs restrict tau spread in astrocytes and neurons

Because phospholipids are transferred to the site of damage as part of repair in the LLOMe paradigm, we next investigated the role of lipid transfer proteins and their upstream regulators. The PITT pathway is initiated by the recruitment of the lipid kinase PI4K2A. PI4K2A locally synthesizes phosphatidylinositol-4-phosphate (PI4P) on the damaged lysosomal surface, driving the formation of expanded membrane contact sites with the endoplasmic reticulum (ER)^18,22^. ATG2A/B, which also known for driving phagophore expansion during macroautophagy by channeling ER lipids^24,25^ and other BLTPs, including VPS13C, operate at these interfaces to further facilitate the bulk lipid transfer required for membrane patching^23^ (Figure 2a).

**Figure 2:**
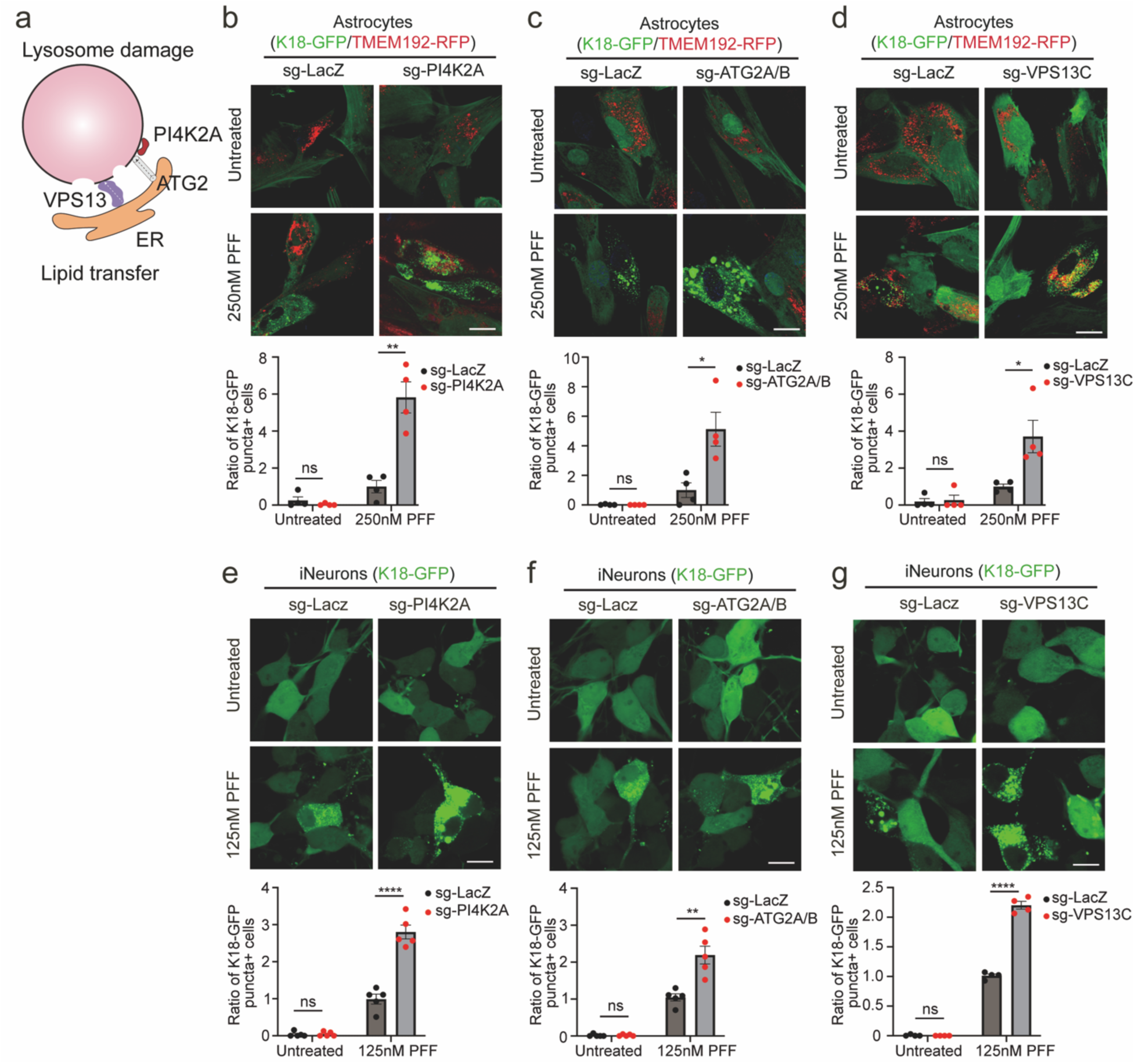
Lipid transport pathways are involved in preventing Tau escape. (a). Diagram of lipid transfer-mediated lysosome repair. (b). K18-GFP/TMEM192-RFP-expressing astrocytes were infected with sg-PI4K2A lentiCRISPR virus, followed by Tau PFF treatment. Then the ratio of K18-GFP puncta-positive cells was quantified (n = 4 independent experiments, 542-667 cells per group were quantified). (c). K18-GFP/TMEM192-RFP-expressing astrocytes were infected with sg-ATG2A/B lentiCRISPR virus, followed by Tau PFF treatment. Then the ratio of K18-GFP puncta-positive cells was quantified (n = 4 independent experiments, 390-522 cells per group were quantified). (d). K18-GFP/TMEM192-RFP-expressing astrocytes were infected with sg-VPS13C lentiCRISPR virus, followed by Tau PFF treatment. Then the ratio of K18-GFP puncta-positive cells was quantified (n = 4 independent experiments, 443-1013 cells per group were quantified). (e). iNeurons were infected with sg-PI4K2A lentiCRISPR virus to achieve PI4K2A knockdown, followed by K18-GFP lentivirus infection and tau PFF treatment. Then the relative ratio of K18-GFP puncta-positive cells was quantified (n = 5 independent experiments, 446-699 cells per group were quantified). (f). iNeurons were infected with sg-ATG2A/B lentiCRISPR virus to achieve ATG2A/B knockdown, followed by K18-GFP lentivirus infection and tau PFF treatment. Then the relative ratio of K18-GFP puncta-positive cells was quantified (n = 5 independent experiments, 524-645 cells per group were quantified). (g). iNeurons were infected with sg-VPS13C lentiCRISPR virus to achieve VPS13C knockdown, followed by K18-GFP lentivirus infection and tau PFF treatment. Then the relative ratio of K18-GFP puncta-positive cells was quantified (n = 4 independent experiments, 164-389 cells per group were quantified). The ratio of K18-GFP puncta+ cells shown in the histogram was normalized to that of the Tau PFF-treated sg-LacZ group. An unpaired two-tailed Student’s t-test was used for pairwise comparisons. Scale bars were 20μm in astrocytes, and 10μm in iNeurons.

To determine whether the mobilization of this ER-to-lysosome lipid transport machinery is required to slow tau escape, we knocked down PI4K2A, ATG2A/B, and VPS13C in the astrocyte reporter cell line. Disruption of any of these lipid transport mediators significantly elevated the ratio of K18-GFP puncta-positive cells following tau PFF treatment (Figure 2b-d). iNeurons depleted from PI4K2A (Figure 2e), ATG2A/B (Figure 2f), or VPS13C (Figure 2g) similarly displayed a pronounced increase in relative K18-GFP aggregation upon PFF treatment.

### Cryo-ET of lysosomes in tau PFF-treated astrocytes

To investigate the structural consequences of lysosomal uptake of tau PFFs, we performed cryo-electron tomography (cryo-ET) on human brain-derived astrocytes expressing our reporter system. We utilized correlative light and electron microscopy (CLEM) to identify target sites positive for AF647-Tau, TMEM192-RFP, and K18-mClover (Figure S3a). To enhance the apparent resolution and contrast of our fluorescence localization, we developed a computational deconvolution pipeline that allowed us to achieve organelle-level precision on our focused ion beam (FIB) milled lamellae (Figure 3a). This pipeline enables the clear identification of compromised lysosomes at tomographic level (Figure 3b), which were verified using the flotillin complex as a membrane identity marker (Figure S3b).

**Figure 3:**
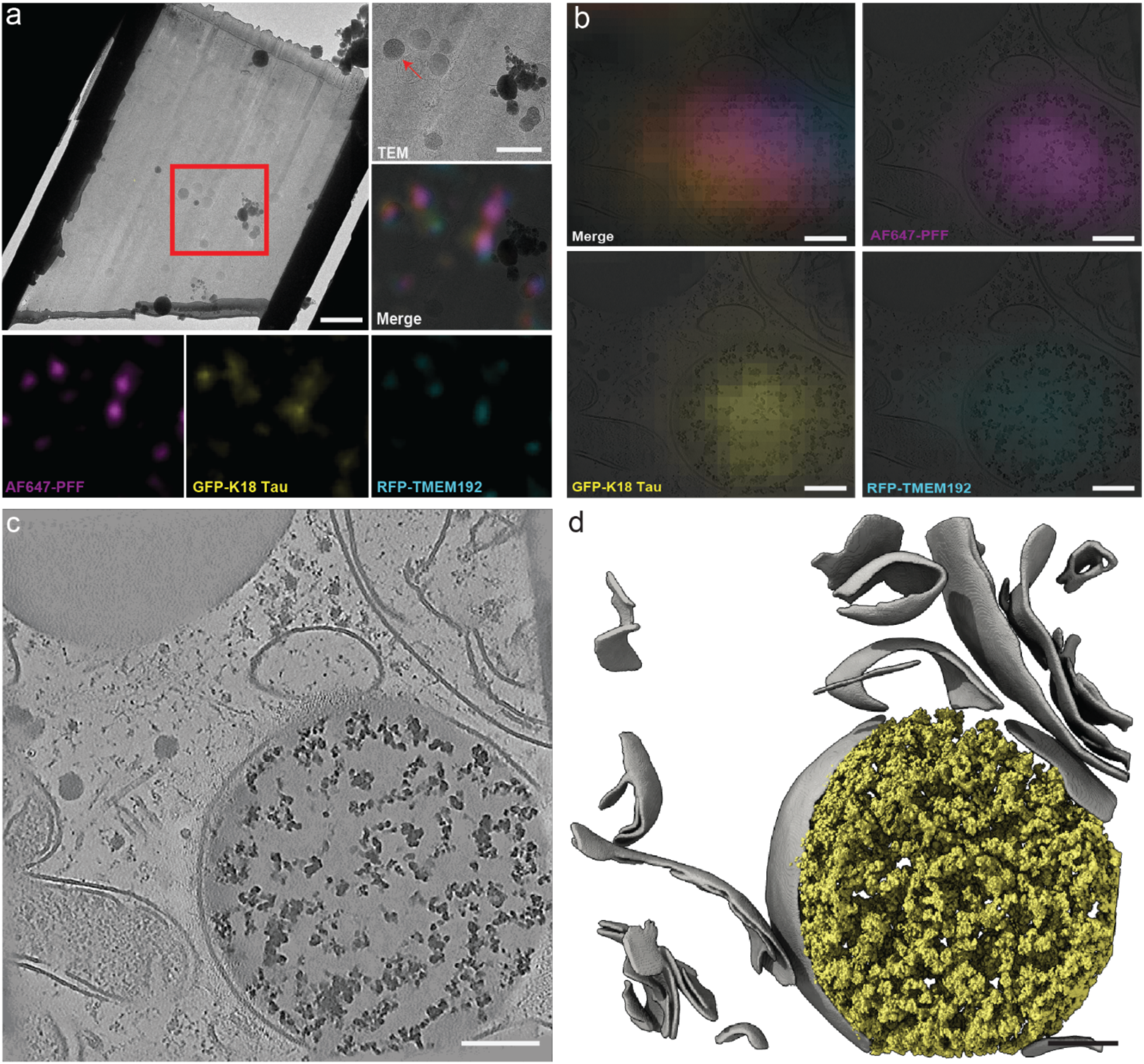
In situ Cryo-ET and computational deconvolution of tau filled lysosomes. (a) Computational deconvolution pipeline for Correlative Light and Electron Microscopy (CLEM) on FIB-milled lamellae, enabling organelle-level precision in localizing tau and RFP-TMEM192 positive compartments. Scale bars are 1.5 um for the lamella and 600 nm medium magnification images (b) Representative slice through a cryo-tomogram and corresponding fluorescence data identifying a compromised lysosome (red arrow in a). Scale bar is 100 nm. (c) 2D tomographic slice showing a highly electron-dense lysosome filled with fragmented tau fibrils after incubation with tau PFFs. Scale bar is 100 nm. (d) 3D segmentation of the lysosomal limiting membrane (gray) and intraluminal tau fragments (yellow). Scale bar is 100 nm.

After incubation with tau PFFs, cryo-ET revealed electron-dense lysosomes filled with fragmented fibrils (Figure 3c). While this dense intraluminal architecture visually resembled the high density of lipid droplets (Figure S3c), these target organelles were unambiguously distinguished by the presence of an intact enclosing lipid bilayer as well as flotillin decoration. This observation suggests that internalized, sonicated tau fibrils undergo partial degradation and condense into densely packed proteinaceous material within the lysosomal lumen. Comprehensive 3D segmentation of both the intraluminal tau fibrils and the limiting lysosomal membrane revealed no instances of fibrils physically piercing or directly protruding the bilayer (Figure 3d, S3d).

### Intralysosomal aggregates in tau PFF-treated astrocytes

Our structural imaging also revealed swollen lysosomes, reaching up to 2 µm in diameter (Figure S3e), that were densely packed with protein aggregates and internalized membrane material (Figure 4a). To visualize the intralumenal aggregates, we segmented the protein fraction and compared voxel counts against the background signal to determine the protein concentration (Figure 4b). Based on a segmented protein volume fraction of 45.0% ± 2.7%, the bulk mass concentration of the aggregate was calculated to be 610 ± 37 mg/mL. This corresponds to an local molarity of 15.0 ± 0.9 mM assuming a composition of pure 0N4R tau or 12.0 ± 0.7 mM assuming an average lysosomal protein population^27^. These measurements align with established densities for pathological aggregates of 600 mg/mL^28,29^, as compared to the normal macromolecular density in the cytosol of 200–300 mg/mL (Figure 4c)^30^.

**Figure 4:**
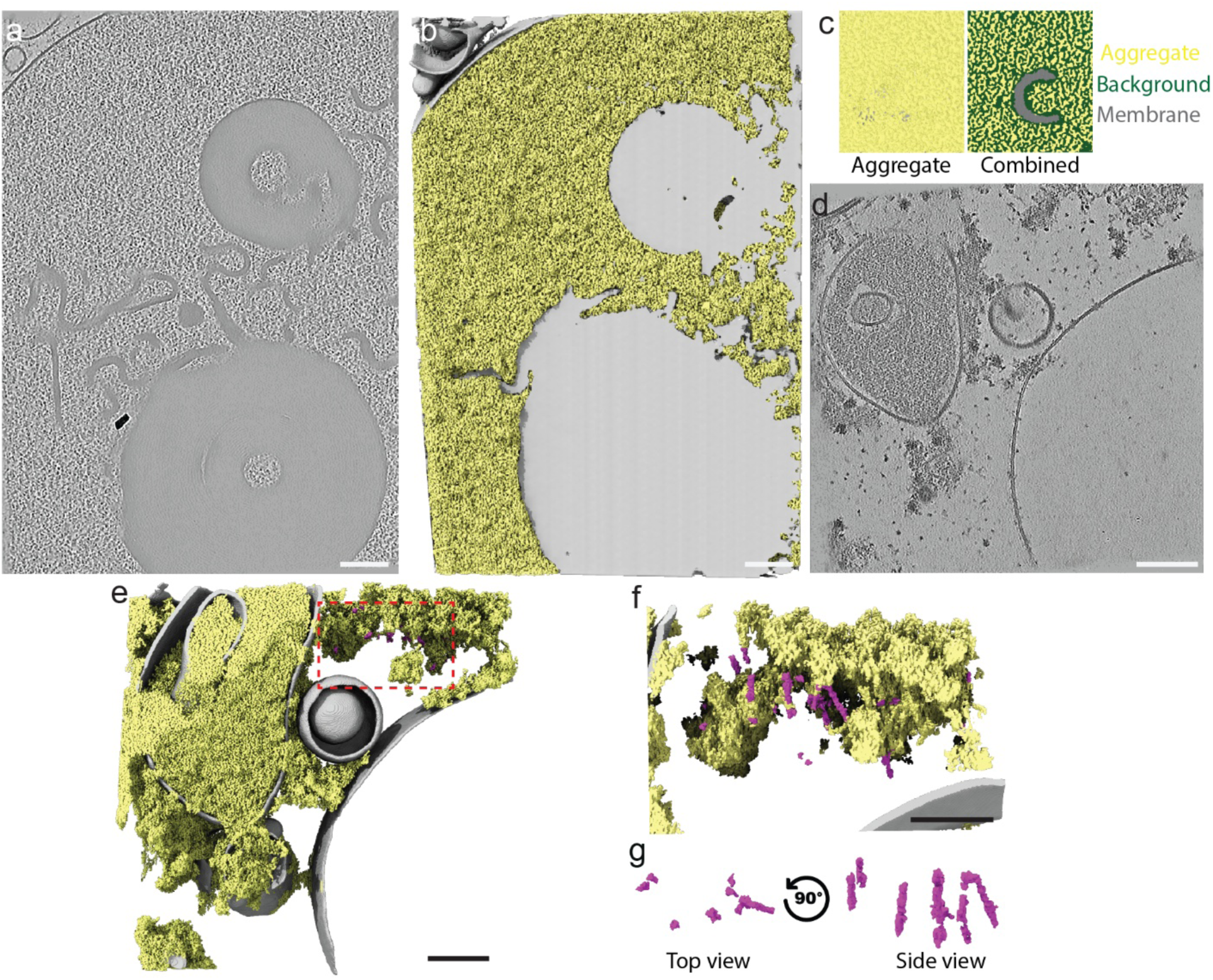
Massive intraluminal tau crowding drives cytosolic escape without macroscopic rupture. (a) Representative cryo-tomogram of a heavily distended lysosome following PFF treatment, showing dense packing of protein aggregates and internalized membrane remnants. (b) 3D segmentation and volumetric analysis of the intraluminal protein fraction (yellow). (c) Exemplary box used for quantification of the intraluminal mass concentration with an average of a segmented volume fraction (44.84% ± 2.68%). (d) Cryo-tomogram captured 48 hours post-PFF treatment, showing escaped cytosolic aggregates and de novo fibril formation in the immediate periphery of a compromised lysosome (e). 3D segmentation of the limiting membrane (gray), the adjacent cytosolic aggregates (yellow) and de novo fibrils (purple). F) Close up view from e showing direct contact of the aggregates and the fibrils. g) Isolated view of the firbils with an average width of 8 nm. All scale bars are 100nm.

Following 48 hours of PFF treatment, we observed cytosolic aggregates positioned directly adjacent to membrane compartments alongside the formation of *de novo* cytosolic fibrils (Figure 4d, SI3e). Comprehensive 3D segmentation of the limiting membranes, the condensed intraluminal aggregates, and the cytosolic fibrils (Figure 4e-g) again revealed no instances of fibrils physically puncturing or mechanically protruding the bilayer.

### Comparative analysis of LLOMe and tau effects on lysosome membranes

To further interrogate the hypothesis of nanoscale versus macroscopic membrane damage, we assessed the endolysosomal limiting membranes for physical hole formation across wild-type (WT), tau PFF-treated, and L-leucyl-L-leucine methyl ester (LLOMe)-treated cells. LLOMe was utilized as a positive control, as it is a well-characterized lysosomotropic agent known to induce large-scale, macroscopic membrane rupture^31^. During our initial tomographic reconstructions, we observed apparent membrane discontinuities or “holes” in the LLOMe treated (Figure 5 a,b) and even within the untreated datasets (Figure SI3 f,g). However, following analysis revealed that in the untreated dataset these gaps consistently aligned with the tilt axis and were often obstructed by adjacent organelles or dense protein assemblies. This strict orientation indicates that these features are missing wedge artifacts inherent to cryo-ET data acquisition, rather than true physical perforations. In contrast, the LLOMe-treated datasets exhibited disrupted lysosomes featuring verifiable physical holes positioned independently of the tilt axis.

**Figure 5.**
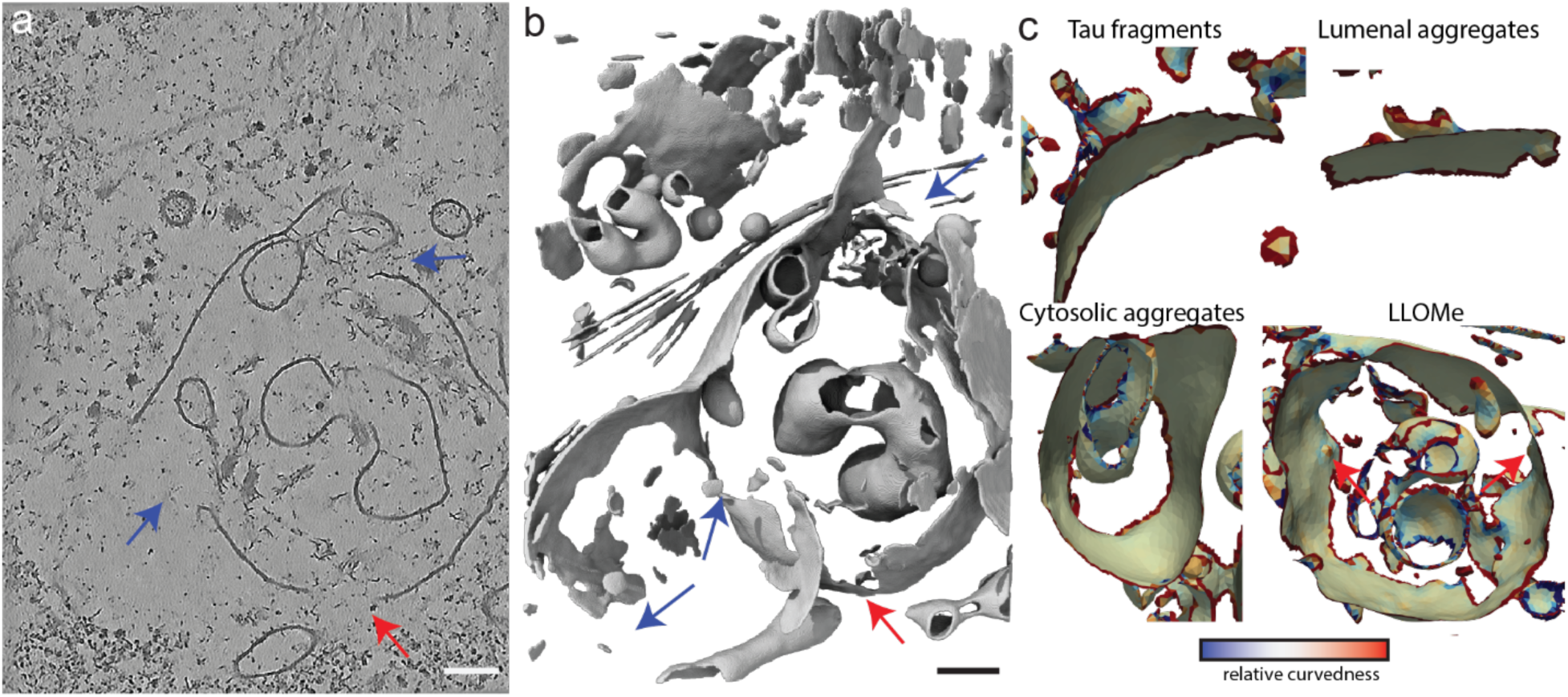
Comparison between tau and LLOMe ehects on lysosomes. (a) Representative tomographic slices of LLOMe damaged lysosome showing apparent membrane discontinuities. Some “holes” (red arrow) align with tilt axis and may be missing wedge artifacts, while other holes are oh the tilt axis and are assigned with confidence (blue arrows). Scale bar is 100 nm. (b) 3D segmentation showing membrane discontinuities (red arrow and blue arrow). Scale bar is 10 0nm. (c) Membrane morphometrics of tomograms shown in 3c, 4a, 4d and 5a showing high local curvature and membrane deformation only on LLOMe treated lysosomes.

To quantitatively compare the nature of membrane stress, we applied a membrane morphometrics pipeline to evaluate local membrane curvature in the tau tomograms (Figures 3D, 4B, and 4F) against the LLOMe-induced damage (Figure 5a). We observed increased local membrane curvature after LLOMe treatment (Figure 5c), but not tau PFF treatment. These data argue against a common mechanism for damage by LLOMe and tau PFFs.

## Discussion

Here we sought to address two outstanding questions in the field. The first is to what extent the host cell cellular repair machinery counteracts tau spread, and the second is the physical mechanism of seeded aggregation in the context of the barrier formed by the lysosomal membrane. With respect to the first question, systematic depletion of ALG-2, CASM, VPS13C, and PITT factors shows that these processes are in fact deployed to protect lysosomes in cells of the CNS, as expected given the precedent of the ESCRT system^13,14^. The disruption of the ALG-2, CASM, or endoplasmic reticulum-to-lysosome bulk lipid transport by PITT exacerbated the cytosolic escape and subsequent aggregation of tau to varying extents in astrocytes and neurons. In iNeurons, knocking down TECPR1 or ATG12 did not promote K18-Tau aggregation, a result likely attributable to the attenuated CASM reported in neurons as compared to other cell types^32^. In contrast, both TECPR1 and ATG12 knockdowns strongly promoted tau seeding in astrocytes. The nearly five-fold increase in seeding seen in the TECPR1 knockdown in astrocytes, with almost every cell manifesting seeding, was one of the strongest sensitizing effects seen. Consistent with the cell type-specific differences seen here, differences in the mobilization of distinct lysosome repair machineries in astrocytes and neurons have also been reported in the context of LLOMe damage^33^. These data support a common underlying mechanism for the repair of LLOMe-induced damage and the suppression of tau spread that operates, albeit with cell type-specific variations in neurons and astrocytes.

Knockdown of the PITT components PI4K2A and ATG2A/B, and another BLTP, VPS13C, led to strong increases in seeding in both astrocytes and neurons. These findings are consistent with the strong role for PITT component PI4K2A in LLOME damage repair in both cell types^33^. The finding of a strong phenotype in neurons is consistent with the finding that VPS13C, the product of an early onset Parkinson’s disease gene^34,35^, is important in neuronal lysosome dynamics^36^. To our knowledge, previous comparative analyses of the function of ATG2A/B in repair in neurons or astrocytes have not been reported. While these data are consistent with a role for ATG2A/B in membrane repair, ATG2A/B, unlike PI4K2A or VPS13C, also has a fundamental role in autophagy^24^. While other interpretations cannot be completely ruled out, these data are most consisted with a tau spread-counteracting role for the PITT pathway augmented by VPS13C in both neurons and astrocytes.

Our data allows us to compare the pathophysiological consequences of tau PFF treatment and the acute damage caused by the experimental lysosomotropic agent LLOMe. LLOMe has long been the tool of choice to study lysosomal membrane damage because of its strong and rapid effects. Recent *in situ* cryo-ET analysis has defined the mechanism of action of LLOMe. Processed by resident cathepsins, LLOMe rapidly condenses into highly ordered amyloid structures^31,37^. These fast-forming LLOMe amyloids possess exposed hydrophobic tips that function as physical wedges, actively perturbing the curvature of the lipid bilayer^31^. The unexpected finding that LLOMe is an amyloid raised the intriguing hypothesis that there could exist a common mechanism for membrane damage shared by synthetic and naturally occurring pathological membrane damaging agents. Moreover, a risk mutant of TMEM106B, when overexpressed, was found to fibrillize within the lysosome lumen in neurons^38^. TMEM106B fibrils were found to deform and rupture membranes in a manner analogous to LLOMe^38^. However, our segmentation and morphometric data, carried out in parallel with LLOMe controls that reproduced the observations of Li et al.^31^, showed no evidence that tau fibrils perturb the curvature of the lysosomal membrane.

We calculated on the basis of segmented ET densities that intralumenal aggregates reached concentrations of up to ∼600 mg/ml over length scales of up to a micron. This concentration value is roughly double the normal density of proteins in the crowded cytosol^30^. Typical *in vitro* tau aggregation and seeding assays rely on concentrations in the range of 0.5 to 5 mg/mL to induce fibrillization^39,40^. It seems difficult to account for these very high concentrations solely on the basis of concentration of endocytosed material. Indeed, these aggregates do not contain features that are recognizable as tau fibrils. These massive aggregates most likely contain additional materials that are incorporated into the aggregate once the aggregation process is initiated by tau PFFs. These data put the mechanism of membrane damage by tau PFFs^14^ into new perspective. The absence of tip-to-membrane interactions and membrane curvature perturbations in the PFF-treated lysosomes argues against the needle-like damage mechanism inferred for LLOMe amyloid. On the other hand, the presence of massive, dense aggregates and lysosomal swelling seen here and previously^14^ suggests that the aggregate itself stresses the membrane, possibly by acting as a surfactant^2^ where it touches the membrane.

Pathological tau exists as distinct structural polymorphs across different tauopathies^41^. Varying fibril architectures possess unique surface hydrophobicities, flexibilities, and packing densities, therefore potentially exerting different stresses on membranes^41,42^. In this study, we could not confidently determine the polymorph of our sonicated and digested tau PFF, nor the *de novo* cytosolic K18 tau fibrils due to heterogeneity induced by heparin and the K18 construct. The average PFF width of 8 nm does at least agree with previous publications^43,44^. Therefore, one limitation of this study is that it is not possible to rule out that other tau polymorphs have a more direct, LLOMe-like, interaction with membranes. We can, however, conclude with confidence that (1) direct membrane disruption is not an inherent, universal property of amyloids, and (2) direct membrane disruption is not necessary for tau PFF seeding and spread.

Tau spread is counteracted by the cellular machinery responsible for repairing nanoscale damage to lysosome membranes, from the ESCRTs^13,14^ to CASM to VPS13C to PITT. This implies that nanoscale pores must open in the course of tau spread. The non-observation of such pores in our study suggests that they are rare, and likely to be transient rather than permanent. While a number of candidate membrane discontinuities were observed in our tomograms, in each case, these apparent ‘holes’ were located in proximity to the tilt axis. It was therefore not possible to unambiguously distinguish these apparent gaps from orientational imaging artifacts^45^. The absence of ‘holes’ from portions of the tomograms far from the tilt axis suggests that they are not abundant in astrocytes. FIB-SEM analysis of neurons has however found constitutive perforations in lysosomal membranes at the rate of a few percent^46^. It was therefore proposed that transient, nanometer-scale pores occur spontaneously in neurons^46^, and that these pores allow cytosolic tau to enter the interior of the lysosome and to polymerize there^47^. The presence of massive aggregates in astrocyte lysosomes seen here suggests that, as for neurons, cytosolic tau can enter the interior of astrocyte lysosomes, thus feeding the aggregates. In contrast to Sanyal et al.^47^, we observed extra-lysosomal aggregation in both neurons and astrocytes. Consistent with our previous finding by STORM microscopy that seeded cytosolic tau aggregates occur proximal to lysosomes in astrocytes, our tomograms showed that PFF-induced aggregates in the cytosol were found adjacent to lysosomes^14^. The appearance and density of these lysosome-proximal cytosolic aggregates was essentially identical to that of the intralysosomal aggregates. The most parsimonious interpretation is that the aggregates originate in a common source, namely, the lysosome itself.

What do these findings imply for therapeutic strategies aimed at reducing tau spread in the CNS? With respect to neurons, defects in PITT components and VPS13C have a stronger effect on seeding than other factors (Figure 2e,g). This seems consistent with the Parkinson’s phenotype of VPS13C mutations^34,35^ and it will be interesting to pursue this point further in the context of dopaminergic neurons and α-synuclein. The molecular functions of VPS13C and ATG2A/B as BLTPs that mediate ER to lysosome phospholipid transfer^17,18^ suggest the potential for boosting ATG2A/B activity or expression to compensate for VPS13C loss of function.

With respect to the larger question of tau escape and seeding, our observations and those of Sanyal *et al*.^47^ suggest a four-way kinetic competition between fibril degradation, cytosolic tau entry, aggregation, and aggregate escape. Strategies that increase the rate of degradation, decrease the rate of aggregation, or decrease the rates of cytosolic tau entry and aggregate escape should be helpful. Chen and Hanson^48^ articulated that lysosome repair factors are not only involved in acute damage response, but also in long-term basal resilience under normal conditions. Strategies that boost these factors should therefore not only help with response to acute damaging agents such as LLOMe, but also with overall lysosome resilience. Higher activity of these factors would be expected to slow the rate of pore formation and slow the entry of cytosolic tau and the egress of aggregate material from the lysosome to the cytosol. A variety of strategies operating at the levels of the ESCRT system, CASM, VPS13C, PITT, or the transcriptional regulation or signaling to these systems could also be contemplated.

## Materials and Methods

### Cell Culture and Cell Line Generation

Human primary astrocytes (ScienCell, 1800) were cultured in ScienCell Astrocyte medium (ScienCell, 1801). iPSCs (NGN2 driven i3Neuron line, kindly provided by Dr. Michael Ward^49^) were maintained on Matrigel (Corning, 354277)-coated plates with Essential-8 Medium (Thermo Fisher Scientific, A1517001). Additionally, Lenti-X 293 cells (utilized for lentivirus production) were cultured for up to 20 passages in DMEM containing 10% fetal bovine serum, Pen/Strep (Life Technologies, 15140122), and L-glutamine (Life Technologies, 25030081).

All cell lines were routinely screened for Mycoplasma contamination and maintained at 37°C with 5% CO₂ in a humidified, copper-lined Heracell VIOS 160i tissue culture incubator (Thermo Fisher Scientific, 51033574).

### Differentiation of iPSCs into iNeurons

iPSCs were differentiated into iNeurons as previously described^50,51^. Briefly, iPSCs were dissociated using StemPro Accutase (Gibco, A1110501) and seeded onto Matrigel-coated plates in induction medium consisting of DMEM/F:12 knockout medium (Thermo Fisher Scientific, 12660012) supplemented with N2 supplement (Thermo Fisher Scientific, 17502048), non-essential amino acids (NEAA; Gibco, 11140050), GlutaMAX (Gibco, 35050-061), and 2 µg/mL doxycycline (Sigma, D9891). Induction medium was refreshed daily for 3 days. Cells were then dissociated and replated onto plates coated with 100 µg/mL poly-L-Ornithine (Sigma, P3655) and 10 µg/mL laminin (R&D Systems, 3446-005-01) in BrainPhys neuronal medium (STEMCELL Technologies, 5790) supplemented with B-27 plus (Gibco, A3582801), 10 ng/mL BDNF (PeproTech, 450-02-10UG), 10 ng/mL NT-3 (PeproTech, 450-03-10UG), 1 µg/mL Laminin, and 2 µg/mL doxycycline. The medium was replaced at half volume every 3 days until cells were harvested for imaging.

### Lentivirus Production

Lenti-X 293 cells (2×10^6) were seeded onto 10 cm dishes and cultured overnight for attachment. Transfection was performed the following day when cells reached approximately 90% confluency. To generate K18.Tau-GFP or TMEM192-RFP lentiviruses, cells were transfected with 6 µg of K18.Tau-GFP (Addgene, 133058) or TMEM192-RFP (Addgene, 134631), 3 µg of VSV-G (Addgene, 8454), and 3 µg of R8.74 (Addgene, 22036). This mixture was combined with 36 µL Mirus LT1 transfection reagent (Mirus Bio, MIR2300) mixed with 1.5 mL Opti-MEM (Thermo Fisher Scientific, 31985062). For LentiCRISPR virus production, cells were transfected with 6 µg of sgRNA plasmid, 3 µg of VSV-G, and 4.5 µg of psPAX2 (Addgene, 12260), along with 36 µL Mirus LT1 transfection reagent.

At 48 h post-transfection, the viral supernatant was collected, clarified by passage through a 0.45 µm syringe filter (Cytiva, 76479-020), and concentrated 10-fold using a Lenti-X concentrator (Takara Bio, 631231). Stable expression pools were later established by titrating the virus concentrate to achieve an expression efficiency approaching 100% across the cell population.

### K18.Tau Aggregation in Astrocytes and iNeurons

Astrocytes were seeded onto 6-well plates and cultured overnight, followed by infection with K18.Tau-GFP virus. The following day, the culture medium was changed, and the cells were subjected to a second round of infection with TMEM192-RFP virus to establish K18.Tau-GFP/TMEM192-RFP-expressing astrocytes. The resulting K18.Tau-GFP/TMEM192-RFP-expressing astrocytes were then replated into 8-chamber confocal dishes (Thermo Fisher Scientific, 155409), allowed to adhere overnight, and passaged once prior to experiments, followed by infection with lentiCRISPR virus. At 48 h post-infection, cells were treated with 250 nM Tau PFF for an additional 48 h. Cells were subsequently fixed with 4% paraformaldehyde (PFA) (diluted from 16% PFA, Electron Microscopy Science, 15710) and imaged using a Nikon A1R confocal microscope with a 60× objective. iPSCs were differentiated into iNeurons as described above. iNeurons were infected with lentiCRISPR virus on day 2 to achieve knockdown of the indicated gene, and with K18.Tau-GFP lentivirus at day 5 to drive K18.Tau expression. Cells were subsequently treated with 125 nM Tau PFF at day 12, and live-cell images were acquired at day 19.

To knock down the indicated genes, the following target sequences were cloned into lentiCRISPR V2 plasmids (Addgene, 52961):

sg-LacZ: 5’-CCCGAATCTCTATCGTGCGG-3’
sg-ALG-2#1: 5’-ACGTCTTCCGCACGTACGAC-3’
sg-ALG-2#2: 5’-CGATCATCCCGGAGTTGTCC-3’
sg-ATG2A#1: 5’-CGGTGAAAAAGTGCGTGGCG-3’
sg-ATG2A#2: 5’-TTATACCGAACATGGCTACA-3’
sg-ATG2B#1: 5’-CCATCAAGAAGAGGGCCTGC-3’
sg-ATG2B#2: 5’-CCATTTGTCCAAGGGGACCT-3’
sg-ATG12#1: 5’-CCAGCAGGTTCCTCTGTTCC-3’
sg-ATG12#2: 5’-GGCTCCTCCGCCATCTTGCT-3’
sg-PI4K2A#1: 5’-CAGCGGAAGCTACTTCGTCA-3’
sg-PI4K2A#2: 5’-GGGTCACTGACCTTTGTACG-3’
sg-TECPR1#1: 5’-ATACACCGACGTCTGCCCCG-3’
sg-TECPR1#2: 5’-CGAGCCGTGTACTTCCGGCA-3’
sg-VPS13C#1: 5’-AACTTACCATCGTCGACCAT-3’
sg-VPS13C#2: 5’-ATCGCCGAAATCAAGATTCA-3’

### Western Blot

To assess knockdown efficiency, total protein was extracted from the indicated cells at 4 days post-infection (astrocytes) or 7 days post-infection (iNeurons) with lentiCRISPR virus. Cell lysates were prepared using 4% sodium dodecyl sulfate (SDS) lysis buffer consisting of 50 mM Tris-HCl (pH 6.8), 100 mM dithiothreitol (DTT), 4% SDS, 12.5% glycerol, and 0.04% bromophenol blue, supplemented with protease inhibitor. Lysates were subsequently denatured by heating at 95°C for 15 min. Proteins were separated using NuPAGE 4%-12% Bis-Tris Gel electrophoresis (Invitrogen, NP0323BOX) and transferred onto membranes (Thermo Fisher Scientific, 88018) for immunoblotting with the indicated primary and secondary antibodies. Chemiluminescent signals were detected with Super Signal West Atto Ultimate Sensitivity Chemiluminescent Substrate (Thermo Fisher Scientific, A38556) and visualized using a ChemiDoc MP imaging system (Bio-Rad). The following antibodies were used: α-Tubulin (Abcam, ab7291), ALG-2 (Proteintech, 12303-1-AP), TECPR1 (CST, 8097), GAPDH (Proteintech, 60004-1-Ig), ATG12 (Cell Signaling Technology, 2010), PI4K2A (Santa Cruz, sc-390026), ATG2A (Proteintech, 23226-1-AP), VPS13C (Proteintech, 28676-1-AP), and ATG2B (Proteintech, 25155-1-AP).

### Immunofluorescence Staining

Day 19 iNeurons cultured in 8-chamber confocal dishes were washed twice with phosphate-buffered saline (PBS) before fixation with 4% PFA for 10 min at room temperature. Cells were subsequently washed three times with PBS and then permeabilized in 0.1% Triton X-100 (in PBS) for 10 minutes. Non-specific binding was blocked by incubating cells in 4% BSA dissolved in 0.05% saponin for 1 h at room temperature. Primary antibody staining was performed by incubation with MAP2 antibody (Proteintech, 17490-1-AP) at 4°C overnight. Cells were washed three times with 0.05% saponin before a 1 h incubation at room temperature with goat anti-rabbit IgG (H+L) highly cross-adsorbed secondary antibody conjugated to Alexa Fluor 647 (Thermo Fisher Scientific, A21245). Following three further washes with 0.05% saponin, nuclei were counterstained with DAPI (0.5 µg/mL). Cells were stored in PBS until imaging, and fluorescence images were acquired using a Nikon A1R confocal microscope using a 60× objective.

### Recombinant 0N4R Tau Protein Purification and PFF Fibrilization

The production of recombinant 0N4R tau protein was carried out following an established protocol^14^, with minor modifications. Briefly, a construct encoding an N-terminally 6× His-tagged human wild-type 0N4R tau was transformed into Rosetta 2 (DE3) competent cells. Transformed bacteria were cultured in LB medium supplemented with ampicillin (100 µg/mL). Upon reaching an optical density of 0.6 at 600 nm (OD₆₀₀), recombinant protein expression was induced by the addition of 1 mM IPTG, and cultures were maintained at 37°C for an additional 3 hours. Bacterial pellets were resuspended and lysed via tip sonication in Buffer A (1x PBS, pH 7.5, containing 2 mM MgCl₂, 10 mM EGTA, 0.5 mM TCEP), to which 0.1 mM PMSF and two EDTA-free protease inhibitor tablets (Thermo Fisher Scientific, PIA32965 / Roche) had been freshly added. Cell debris was removed by centrifugation, and the clarified lysate was loaded onto Pierce High-Capacity Ni-IMAC Resin (Thermo Fisher Scientific, A50585). His-tagged tau was selectively retained on the resin and subsequently released by competitive elution using Buffer A supplemented with 500 mM Imidazole (pH 7.5).

His-tag cleavage was performed by adding TEV protease directly to the IMAC eluate, followed by extensive dialysis at 4°C overnight against Buffer B (20 mM MES, pH 6.8, 50 mM NaCl, 1 mM EGTA, 1 mM MgCl₂, 0.5 mM TCEP). Ion exchange chromatography was then performed on a 5 mL HiTrap SP column (Cytiva) to further purify tau from contaminants, employing a 0–1 M NaCl linear gradient for elution. Peak fractions containing tau, as confirmed by Coomassie blue staining of SDS-PAGE gels, were pooled and subjected to size exclusion chromatography on a Superdex 75 16/60 column (Cytiva) pre-equilibrated with SEC buffer (1x PBS, pH 7.5, 2 mM MgCl₂, 1 mM TCEP). After determining the concentration via absorbance at 280 nm, the purified protein was concentrated to approximately 20 µM, snap-frozen in liquid nitrogen, and stored in 75 µL aliquots at −80°C until use.

To generate tau PFF, purified monomeric tau (10 µM) was mixed with heparin (10 µM; Thermo Scientific, cat# A16198.03) and incubated at 37°C with continuous agitation at 1,200 rpm overnight to promote fibril nucleation and elongation. The structural integrity of the resulting fibrils was verified by negative-stain transmission electron microscopy (TEM) or cryo electron microscopy. The resulting fibrils were then fluorescently labeled using an AlexaFluor 647 NHS-ester dye (Invitrogen, A20006) in accordance with the manufacturer’s instructions. Prior to dye addition, the pH of the mixture was raised by adding NaHCO₃ to a final concentration of 100 µM. The dye was introduced at a 3:1 molar ratio of monomeric protein to dye. Following a 1-hour incubation in the dark at room temperature, any remaining unlabeled dye was removed using a Zeba 7k MWCO spin desalting column (Thermo Fisher Scientific, 89882). Prior to use in cell-based seeding assays, fibrils were fragmented into short seeds by tip sonication using a pulsed program (1s on / 1s off) for a total of 60s.

### Data Analysis

All statistical analyses were conducted using GraphPad Prism 10. Quantitative data are presented as means ± SEM. An unpaired two-tailed Student’s t-test was used in this study for pairwise comparisons. Statistical significance was defined as *P* < 0.05, with significance levels indicated as \**P* < 0.05, \*\**P* < 0.01, \*\*\**P* < 0.001, and \*\*\*\**P* < 0.0001; non-significant differences are denoted as “ns”.

### EM grid seeding and cryo-FIB milling

Gold Quantifoil R2/2 200-mesh EM grids (EMS, catalog #Q250-AR2) were glow-discharged for 30 seconds at 15 mA and subsequently floated on drops of 0.01% poly-D-lysine (Sigma, catalog #A-005-M) in a laminar flow hood for 30 minutes. Concurrently, 8-chamber slides with removable wells (Sigma, catalog #PEZGS0816) were coated with 250 µL of 0.01% poly-D-lysine. Astrocytes were split using 0.025% trypsin (ScienCell, 0183) and resuspended to a concentration of 200,000 cells/mL. The poly-D-lysine was aspirated from the chamber wells and replaced with 200 µL fresh medium. The EM grids were rinsed in medium and placed at the bottom of the wells, after which 200 µL of the cell suspension was added atop each grid to achieve a seeding density of 40,000 cells per grid. A day after seeding, cells were treated with 250nM PFF for 24hrs. On the following day, cells were split onto EM grids such that fibril incubation time before freezing was 48hrs and 24hrs on grids to minimize cell stress. This same seeding protocol was used for U2OS cells but at a density of 20k cells per grid and with DMEM. 250uM LLOMe was added to the media 20min prior to plunge freezing. For plunge freezing, a Vitrobot Mark IV (ThermoFisher) was equipped with blotting paper and a Teflon sheet and equilibrated to 37 °C at 90% humidity. The removable wells were detached from the slide, and the grids were retrieved using Vitrobot tweezers. Each grid was washed three times with drops of PBS, back-blotted using a blot force of 10 for 8 seconds and directly plunge-frozen in liquid ethane. The vitrified grids were then clipped into notched bases designed for cryo-FIB milling (ThermoFisher).

The clipped grids were loaded into an Aquilos 2 dual-beam system equipped with integrated fluorescence light microscopy (iFLM, ThermoFisher). Initial screening to identify potential lamellae sites was performed using scanning electron microscopy (SEM). The cells were then iteratively screened using iFLM to precisely target regions exhibiting clustered GFP, RFP and AF647 signals for lamella placement. To protect the sample during milling, the grids were sputter-coated for 15 seconds at 30 mA and 10 Pa, followed by a 1-minute gas injection system (GIS) coating. Automated focused ion beam milling (Auto-TEM) was utilized to generate lamellae with a final thickness of approximately 180 nm at each targeted site. The ablation process was conducted in progressive steps: 0.5 nA of current was used to rough-mill the cellular material down to 3 µm in thickness, followed by 100 pA to thin the sections to 1 µm, 50 pA to reach 500 nm, and finally a 30 pA polishing current to achieve the target 180 nm thickness.

### Cryo-electron tomography data acquisition

Grids containing lamellae were retrieved from the Aquilos and immediately stored in nitrogen or loaded into a 300 kV Titan G3 using a Quantum K3 direct electron detector (Gatan) and images acquired in EFTEM mode with a 25 eV slit width or on a G4 equipped with a cold field emission gun (CFEG), a Selectris X Energy Filter, and a Falcon 4i direct electron detector (Thermo Fisher Scientific, Hillsboro, OR, USA). Those images were acquired in EFTEM mode with a 10eV slit width. The autogrids containing lamellae were loaded such that the pre-tilt axis created by FIB milling was perpendicular to the tilt axis of the microscope. Montage maps were generated for the entire autogrid to identify lamellae positions and a second medium magnification montage generated at each lamellae site. Polygon montages were used to outline the borders of each lamella and used to guide data collection. Tilt-series were collected using PACE-tomo starting from 9-degree lamellae pre-tilt with increments of 3 degrees^52^. The nominal defocus was varied between tilt-series from −2 to −6 μm with a step size of 0.25 μm. The total dose per tilt series was approximately 120 e-/ Å2. Frames were saved in Electron Event Representation (EER) format for Falcon data.

### CLEM data processing

The detailed protocol can be found here: https://dx.doi.org/10.17504/protocols.io.yxmvm8149g3p/v153. Briefly, Fluorescence Z-stacks acquired from the integrated FIB-mill fluorescence module (e.g., Aquilos 2) were processed using Fiji/ImageJ. To improve signal-to-noise ratios and feature resolution, synthetic 3D point spread functions (PSFs) were generated and applied to the fluorescence stacks using Richardson-Lucy 3D deconvolution. The deconvolved stacks were subsequently rotated to correct for specimen tilt. Corresponding brightfield Z-stacks were center-cropped to match the fluorescence channel dimensions and converted to 32-bit format. Finally, all processed channels were merged into a single composite hyperstack to facilitate downstream correlation and manually corelated to the TEM images.

### Cryo-electron tomography data processing and segmentation

Scipion3 was used to facilitate all downstream data processing^54^. Briefly, MotionCor3 was used to motion correct tilt series and binning in Fourier space to the physical pixel resolution was applied during correction^55^. CTFfind5 was used for CTF estimation and AreTomo2 was used for tilt series alignment and tomogram reconstruction^31,56^. Tomograms were denoised using IsoNet2 before using TOMO3D^57,58^. Resulting tomograms were segmented using Membrain with default parameters and with Comet Technologies *Dragonfly 3D World* (Version 2025.1) using the U-Net 3D architecture^59^. Tomograms were visualized using 3DMOD and segmentations using ChimeraX^60,61^. Segmentation with Membrain were used to run the membrane morphometrics with the same settings as previously described^31,62^.

### Calculation of aggregate concentration

Segmentations of the aggregates and background were generated using Dragonfly software. To quantify the volume fraction, bounding boxes were placed within three distinct regions across three separate tomograms, and voxel counts were extracted using the intersection tool. The aggregate volume fraction was calculated for each tomogram and subsequently averaged. The bulk mass concentration of the lysosomal aggregate was derived from the volumetric segmentation using the standard partial specific volume of folded proteins. The segmented protein volume fraction (*Vf*) was determined to be 44.84% ± 2.68%. Assuming an average partial specific volume (*v*) of 0.73 cm³/g for typical globular proteins^63^, the theoretical density of the pure, anhydrous protein (*ρ*) was calculated as the reciprocal of the partial specific volume:

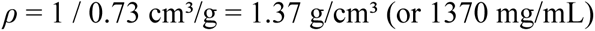

The mass concentration (*Cmass*) of the protein within the segmented lysosomal volume was calculated as the product of the volume fraction and the pure protein density. The standard deviation of the volume fraction was propagated linearly:

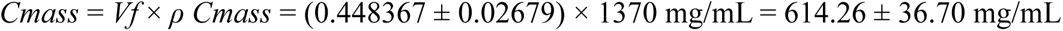

To convert the mass concentration into a molar concentration (*CTau*) assuming the aggregate is composed entirely of Tau 0N4R, the molecular weight (*MW*) of the monomer was utilized (*MWTau* = 42.1 kDa, or 42,100 g/mol).

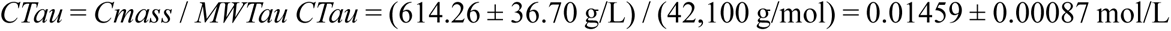

This corresponds to an estimated local molarity of 14.59 ± 0.87 mM for pure Tau 0N4R.

To calculate the collective molar concentration (*CLyso*) assuming the volume consists of a mixed population of typical lysosomal proteins, an adjusted average molecular weight was applied (estimated *MWLyso* = 50.0 kDa, or 50,000 g/mol).

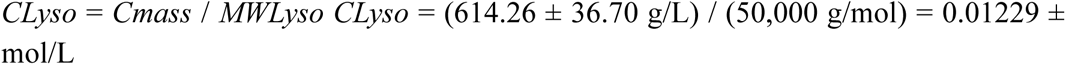

This corresponds to a collective estimated molarity of 12.29 ± 0.73 mM for an average lysosomal protein population.

### Cell lines

Human Osteosarcoma (RRID:CVCL_0042) cells were received from the UCB Cell Culture Facility. U2OS were treated with fresh DMEM media with 10% FBS for 30 minutes prior to being subjected to 250 μM LLOMe for 20 minutes. After incubation, the cells were frozen and subjected to the same tomography workflow as described above.

### Abbreviations

Here is the fully combined and updated abbreviation list, merged with your old list and formatted as a clear text list instead of a table. It is organized alphabetically for easy integration into your manuscript.

## Supporting information

key resource table

## Abbreviations

0N4R: 0 N-terminal inserts, 4 microtubule binding repeats tau isoform
AD: Alzheimer’s disease
AF647: Alexa Fluor 647
ALG-2: Apoptosis-linked gene 2
ALS: Amyotrophic lateral sclerosis
ATG12: Autophagy-related 12
ATG2A/B: Autophagy-related 2A/B
ATG5: Autophagy-related 5
ATG8: Autophagy-related 8
BLTP: Bulk lipid transfer protein
CASM: Conjugation of ATG8s to single membranes
CLEM: Correlative light and electron microscopy
CNS: Central nervous system
CRISPR: Clustered regularly interspaced short palindromic repeats
Cryo-ET: Cryo-electron tomography
DKO: Double knockdown
DMEM: Dulbecco’s Modified Eagle Medium
ELN: Endolysosomal network
ER: Endoplasmic reticulum
ESCRT: Endosomal sorting complexes required for transport
FIB: Focused ion beam
FTD: Frontotemporal degeneration
GFP: Green fluorescent protein
GPN: Glycyl-L-phenylalanine 2-naphthylamide
iPSC: Induced pluripotent stem cell
K18: Tau fragment containing four microtubule-binding repeats
LLOMe: L-leucyl-L-leucine methyl ester
PD: Parkinson’s disease
PFF: Pre-formed fibrils
PI4K2A: Phosphatidylinositol 4-kinase type 2 alpha
PI4P: Phosphatidylinositol-4-phosphate
PITT: Phosphoinositide-initiated membrane tethering and lipid transport
RFP: Red fluorescent protein
TECPR1: Tectonin beta-propeller repeat containing 1
TMEM106B: Transmembrane protein 106B
TMEM192: Transmembrane protein 192
VPS13C: Vacuolar protein sorting 13 homolog C
WT: Wild-type

## Acknowledgements

We thank Pei-I “Leo” Tsai for program and open access compliance support. Portions of this work were performed at the Stanford-SLAC Cryo-ET Specimen Preparation Center, supported by the NIH Common Fund’s Transformative High-Resolution Cryoelectron Microscopy program (U24GM139166); we thank Lydia-Marie Joubert for her assistance. Some of this work was performed at the Midwest Center for Cryo-ET (MCCET) and the Cryo-EM Research Center located in the Department of Biochemistry at the University of Wisconsin-Madison, supported by the NIH Common Fund Transformative High Resolution Cryo-Electron Microscopy program (U24 GM139168), where we thank Anil Kumar for his help correlating iFLM data for tomography data collection.

## Data availability

The data, code, protocols, and key lab materials used and generated in this study are listed in a Key Resource Table alongside their persistent identifiers.

The raw tilt series and a few selected high-quality tomograms were deposited at the Electron Microscopy Public Image Archive (EMPIAR) under the identifier (XXXXXXX). All raw data generated during this study can be found on Zenodo (XXXXXX).

## Declarations

### Ethics approval and consent to participate

Not applicable.

### Consent for publication

Not applicable.

## Supplementary information

**Figure S1:**
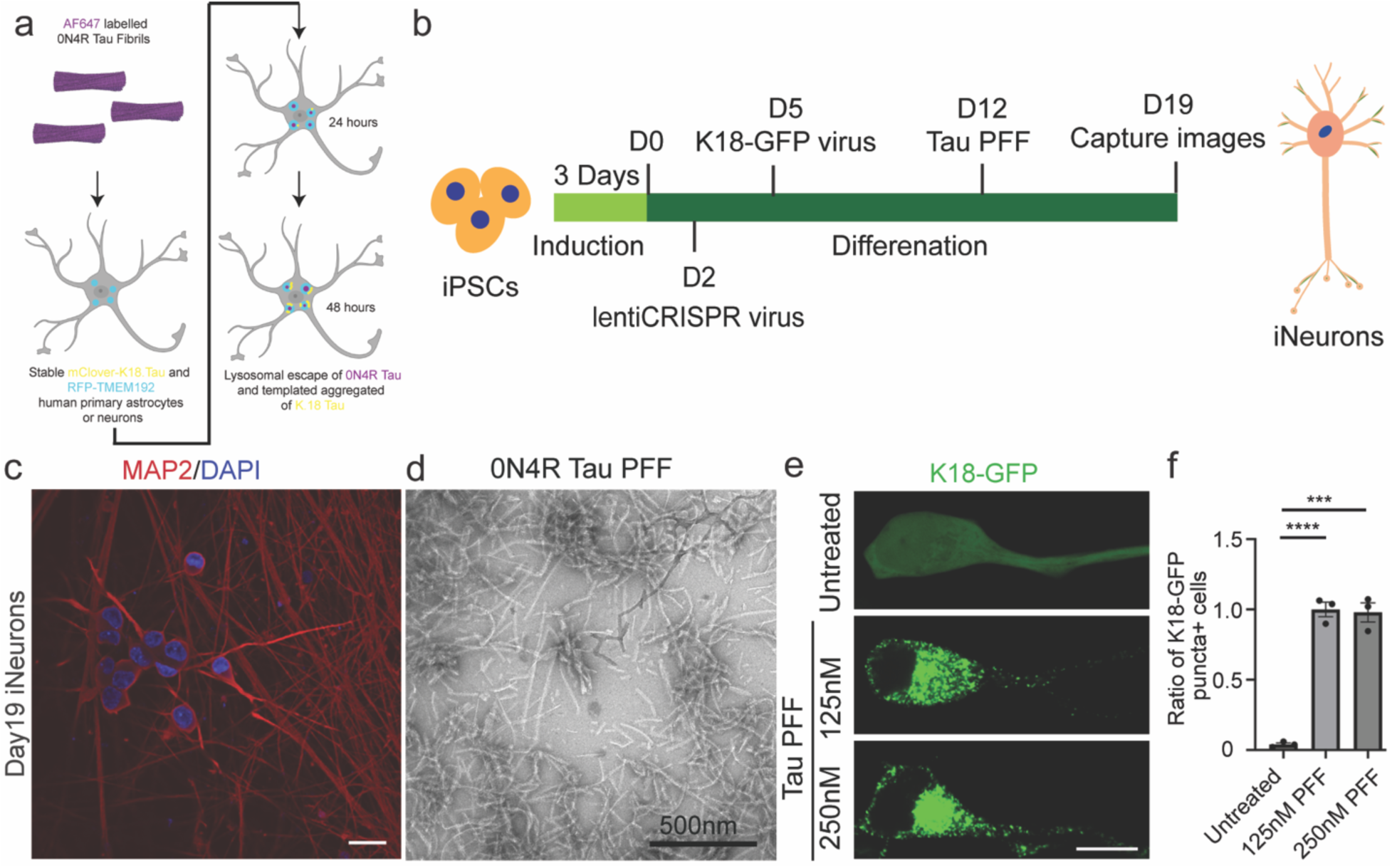
Establishment of pathogenic templating in iPSC-derived human neurons. (a) Schematic of the K18-GFP/TMEM192-RFP reporter system used to monitor lysosomal breach. (b) Experimental timeline for the maturation of iNeurons and subsequent tau PFF treatment. (c) Immunofluorescence characterization of Day 19 iNeurons positive for the mature neuronal marker MAP2. (d) Negative stain images of Tau PFF prior to sonication (e) Comparison of K18.Tau aggregation following exposure to 125 nM and 250 nM tau PFFs, showing successful induction of fluorescent puncta. (f) Evaluation of (e) showing the ratio of k18.tau puncta per cells. The ratio of K18-GFP puncta+ cells shown in the histogram was normalized to that of the 125nM PFF-treated group. An unpaired two-tailed Student’s t-test was used for pairwise comparisons. Scale bars are 20μm.

**Figure S2:**
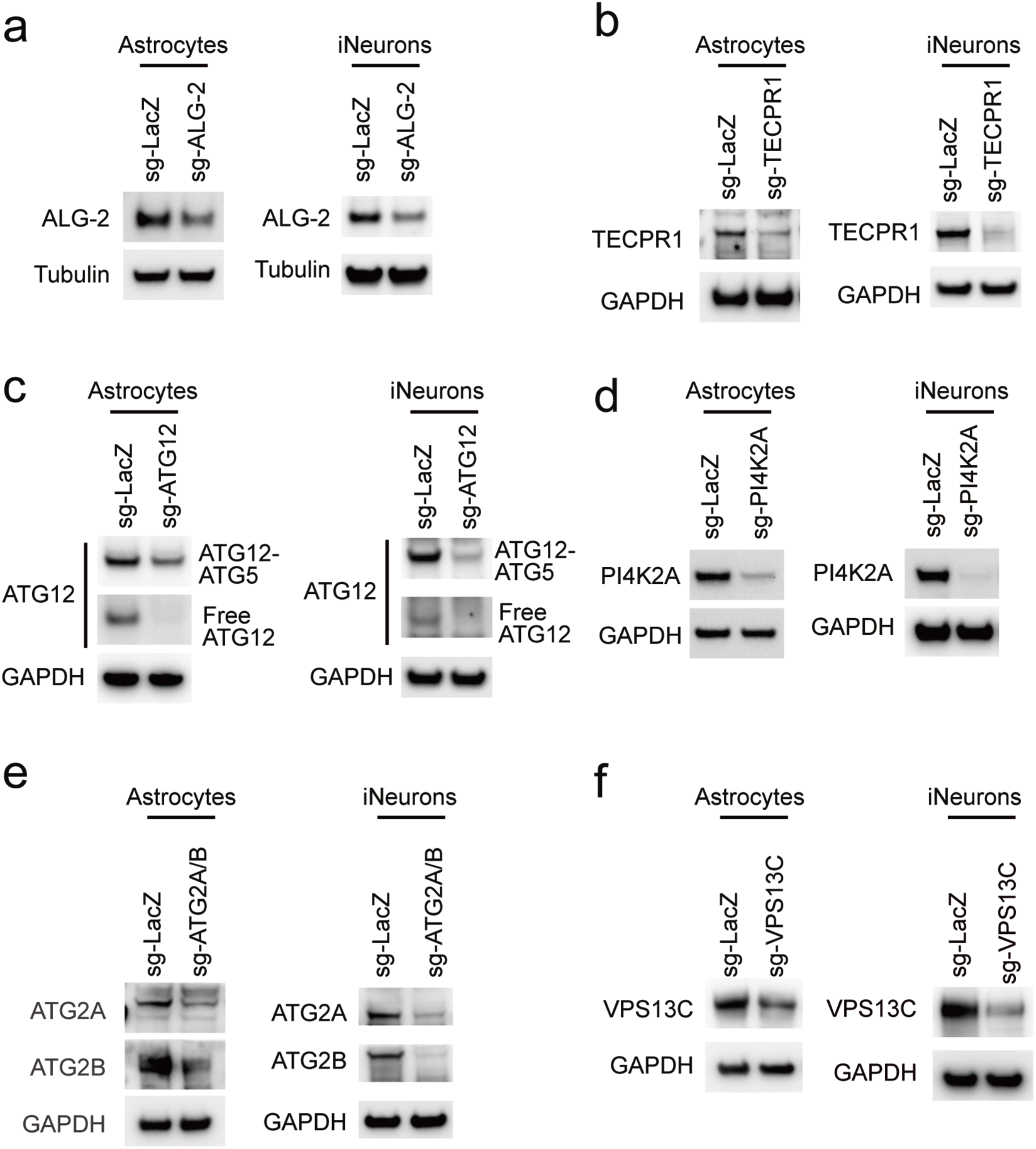
Knockdown ehiciency of the indicated genes in astrocytes and iNeurons. (A). Knockdown ehiciency of ALG-2 in astrocytes and iNeurons was assessed by Western blot. (B). Knockdown ehiciency of TECPR1 in astrocytes and iNeurons was assessed by Western blot. (C). Knockdown ehiciency of ATG12 in astrocytes was assessed by Western blot. (D). Knockdown ehiciency of PI4K2A in astrocytes and iNeurons was assessed by Western blot. (E). Knockdown ehiciency of ATG2A and ATG2B in astrocytes and iNeurons was assessed by Western blot. (F). Knockdown ehiciency of VPS13C in astrocytes and iNeurons was assessed by Western blot.

**Figure S3:**
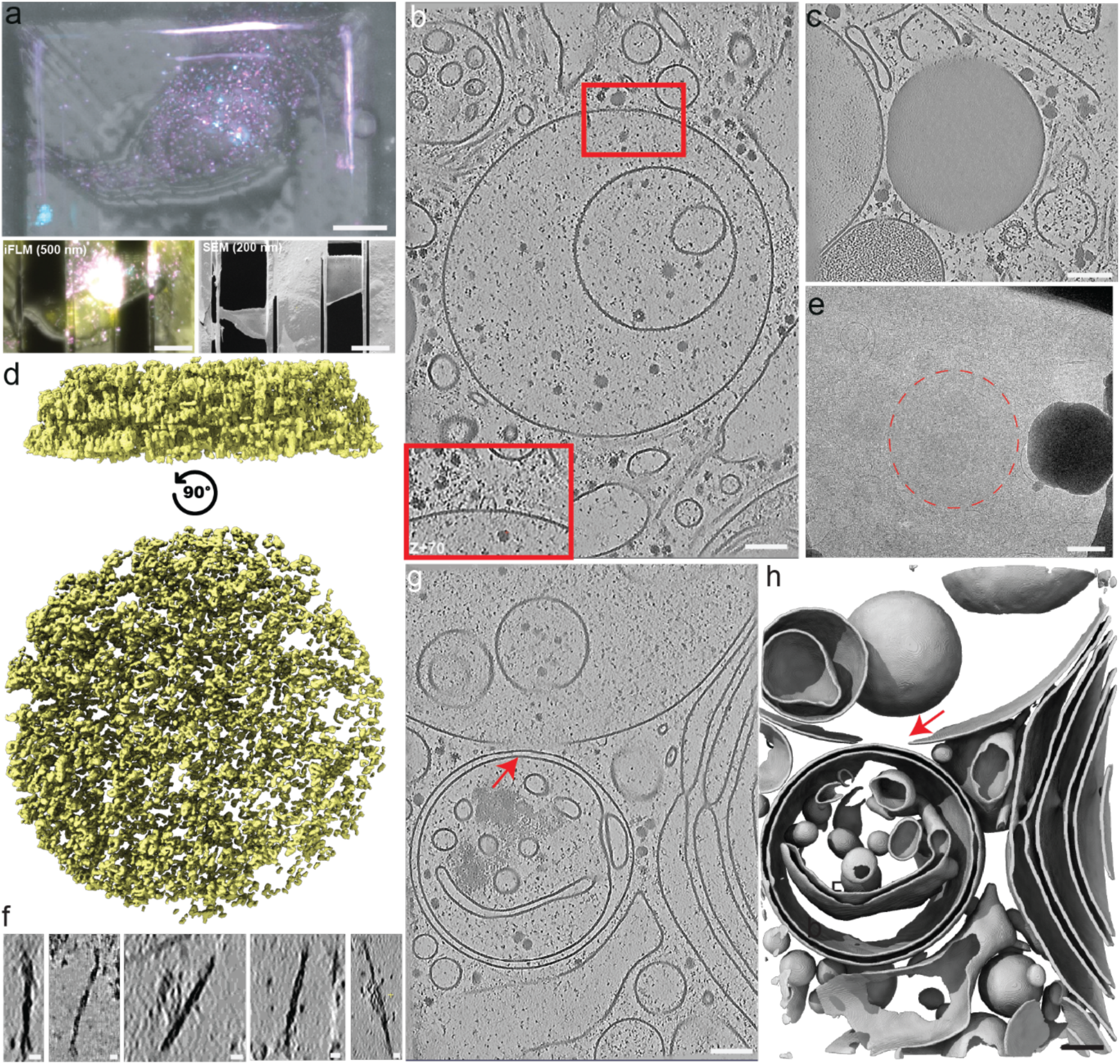
Correlative Validation of Lysosomes. (a) Representative correlative light microscopy (CLEM) images of FIB-milled lamellae. iFLM images were acquired at 500nm and afterwards milled down to 200 nm thickness. Scale bars are 20 and 10 um. (b) Tomographic slice showing the limiting membrane of a target organelle decorated with flotillin complexes (red box), verifying its identity as a lysosome. Scale bar is 100 nm. (c) Structural comparison between a tau-filled lysosome and a cytoplasmic lipid droplet. Sclae bar is 100 um. (d) Additional gallery of 3D segmented intraluminal tau fragments (yellow) viewed from multiple orientations. (e) Medium magnification view of the full swollen lysosome shown in 4a. Sclae bar is 500 nm. (f) Gallery of cytosolic tau fibrils shown in 5f. scale bar is 10 nm. Representative view of a untreated lysosome with a membrane discontinuity (red arrow) aligning with the tilt axis. Scale bar is 100 nm. (h) Membrane Segmentation of f. Scale bar is 100 nm.

